# Individual alpha frequency tACS reduces functional connectivity across the default mode network

**DOI:** 10.1101/2024.08.04.606325

**Authors:** Martín Carrasco-Gómez, Alejandra García-Colomo, Jesús Cabrera-Álvarez, Alberto del Cerro-León, Carlos J. Gómez-Ariza, Andrés Santos, Fernando Maestú

## Abstract

**Objectives:** Research on the influence of transcranial alternating current stimulation over alpha functional connectivity (FC) is scarce, and at the same time poses as a potential treatment for various diseases. This study aimed to investigate the effects of individualized alpha frequency tACS (IAF-tACS) on FC within the default mode network (DMN) in healthy individuals, particularly focusing on the precuneus (PCU) as a major hub within the network.

**Materials and Methods:** 27 healthy participants were recruited, which underwent a 20-minute IAF-tACS session and three magnetoencephalography (MEG) recordings: two pre-stimulation and one post-stimulation. Participants were randomly assigned to either the stimulation or sham group. FC was evaluated through the corrected imaginary phase locking value (ci-PLV) and leakage corrected amplitude envelope correlation (AEC-c). Statistical analyses compared both Pre-Post FC ratio between groups through ratio t-tests and intragroup FC changes through repeated measures t-tests, with FDR correction applied to account for multiple comparisons. An additional analysis simulated the influence of the cortical folding on the effect of tACS over FC.

**Results:** IAF-tACS significantly decreased AEC-c within the PCU and DMN in the stimulation group compared to the sham group, especially influencing antero-posterior links between hubs of the DMN. No significant changes were observed in ci-PLV connectivity metrics. Negative correlations were found between AEC-c FC changes and power alterations in posterior DMN areas, suggesting a complex interaction between cortical folding and electric field direction.

**Conclusions:** Against our initial hypothesis, IAF-tACS reduced FC in the DMN, possibly due to phase disparities introduced by cortical gyrification. These findings suggest that tACS might modulate FC in a more complex manner than previously thought, highlighting the need for further research into the personalized application of neuromodulation techniques, as well as its potential therapeutic implications for conditions like Alzheimer’s disease.

## Introduction

The brain shows spatio-temporally organized patterns of activity even when at wakeful rest. During this resting state, the alpha rhythm (8 – 12 Hz) becomes the predominant oscillation over posterior regions^1^. The spontaneous emergence of alpha in resting state, and its correlation with performance, has led to the hypothesis that alpha reflects a functional inhibitory role that is key to the allocation of processing resources for environmental stimuli^2^. In addition to the alpha rhythm, resting state is associated with changes in functional connectivity (FC) understood as statistical relationships between brain signals over time^3,4^. These functional interactions give rise to brain networks that reflect interregional communication to support cognitive function. The default mode network (DMN) is a key functional network that is active during resting periods and deactivates during task performance^5^. The DMN is involved in memory processing^6^ and includes several regions such as the medial prefrontal cortex, middle temporal gyrus, the hippocampus, the posterior cingulate cortex (PCC) and the precuneus (PCU), the latter standing out as a hub of the network^7,8^ Disruptions in both the alpha rhythms and the DMN have been associated with multiple brain diseases^9–11^, therefore, their modulation constitutes a potential therapeutic avenue to explore.

Transcranial alternating current stimulation (tACS) is a non-invasive stimulation technique capable of modulating oscillatory activity in the brain and inducing neuroplasticity^12^. Weak alternating electrical currents are applied to the scalp, reaching the brain, influencing neurons’ membrane potentials and thus, influencing neuronal probability of activation^13^. As a result, the application of tACS can induce neural entrainment (i.e. the synchronization of an oscillatory system to an external driver) at the frequency of stimulation^14^. The anatomical and functional complexity of the brain makes the results of neuromodulation highly dependent on the protocol employed^15^, but it has been observed that entrainment is more likely to occur if the stimulation frequency matches the frequency of the ongoing oscillation^16^ and if the anatomy of the target region is taken into account^17,18^.

Externally applied oscillatory currents can interact with the endogenous alpha rhythm to enhance or disrupt it^14,19^. In this line, tACS at the individual alpha frequency (IAF-tACS) has been shown to increase the power of alpha rhythm after stimulation with effects remaining up to 70 minutes after the protocol^20,21^. These after-effects likely involve plastic changes in neuronal circuits^22^.

Beyond these effects, cortical modulation may also influence FC. Recent research suggests that tACS modulates FC in a frequency- and network-specific manner. For instance, occipito-parietal alpha stimulation during resting-state increases FC specifically in the alpha band and in the regions of the DMN^23,24^, which were suggested in the latter to be mediated by changes in alpha power induced by stimulation^24^. In contrast, a 6 Hz stimulation in resting state - when the brain is typically oscillating in alpha - decreases FC in the DMN^25^, highlighting the variability in tACS effects depending on the stimulation frequency. These modulations of FC have been shown to enhance cognitive performance increasing interregional communication through coherence^26–30^. Therefore, tACS has become a promising tool for modulating FC both in health and disease.

In this study, we aimed to investigate the influence of a personalized IAF-tACS protocol on healthy individuals’ FC through MEG recordings. According to the aforementioned studies, we expected to find an increased FC in the participants’ DMN after the stimulation. To get a spatially finer analysis, we evaluate the FC of the PCU as one of the main hubs in the DMN. We implement two types of FC metrics to investigate both phase and amplitude connectivity parameters. Finally, we relate changes in FC to changes in power in the areas of interest.

## Methods

### Study design

The study comprised one tACS session and three magnetoencephalography (MEG) scans, two before (*Pre1* and *Pre2* sessions) and one after the stimulation (*Post* session). We firstly recorded the electrophysiological activity of each participant through two successive 5-min eyes-closed resting-state MEG recordings, with a 10-minute interval between sessions. Two MEG recordings before the stimulation were included to account for any possible variability in the individual alpha-peak frequency (IAF) of the participants. The IAF was derived from each recording using a fast-preprocessing algorithm described in the “*MEG preprocessing and source reconstruction*” section, and then averaged. Statistical analyses for the current paper were carried out using the *Pre2* data as the baseline measure since it was closer in time to the stimulation and all participants had experienced a similar situation until that point. Participants were then randomly assigned to either the real stimulation (*stim*) or placebo (*sham*) group, and received 20 min of stimulation at their own IAF with their eyes closed. Immediately after the stimulation, we performed a third MEG recording to measure its effects on the participants. We decided to use a resting-state eyes-closed paradigm given its prominent alpha activity, and previous work on the same kind of stimulation^31,32^.

### Participants

We recruited 27 (11 female) participants aged between 22 and 55 years from the Center for Cognitive and Computational Neuroscience (C3N) at the Complutense University of Madrid (UCM). The study included only right-handed, native Spanish-speaking participants without any prior neuropsychiatric history or metallic prostheses that could interfere with neuroimaging and neuromodulation. Additionally, participants with indistinguishable Individual Alpha Frequency (IAF) were excluded. We adhered to current guidelines and safety regulations throughout the research and obtained informed consent from each participant before their involvement.

### MEG Data acquisition

MEG signals were acquired during 5 min of eyes-closed resting state at 1 kHz sampling rate, using 306 channels (102 magnetometers and 204 gradiometers) whole-head Elekta Neuromag system (Elekta AB, Stockholm, Sweden) located in a magnetically isolated room (VacuumSchmelze GmbH, Hanau, Germany). Using a Fastrak 3D digitizer (Polhemus, Colchester, Vermont), the positions of four head position indicator (HPI) coils attached to the scalp were defined and the shape of each participant’s head relative to three anatomical locations (nasion and both preauricular points) was modeled. An online anti-aliasing filter [0.1–330) Hz] was applied during the whole session.

### MEG preprocessing and source reconstruction

The Maxfilter software (v.2.2, correlation threshold = 0.9, time window = 10 s) was used to remove the environmental noise in the raw data, using the temporal extension of the signal space separation (tSSS) method with movement compensation^33^. For further analysis, only data from magnetometers was considered, given the redundancy in gradiometer data after tSSS^34^. Physiological and jump artifacts were located using the automatic Fieldtrip software^35^ and later reviewed by MEG signal experts. Lastly, the clean signal was divided into 4 second-segments and eye-blinks and cardiac artifacts were removed with an independent component analysis (ICA) based on SOBI^36^. All segments with physiological artifacts were discarded. Preprocessed MEG data was then used to carry out source localization using a Linearly Constrained Minimum Variance (LCMV) beamformer^37^. Since not all subjects had available T1 MRIs, a 1 mm resolution template of healthy adults normalized to the Montreal Neurological institute (MNI) with a 1 mm voxel size template was used to place the sources inside the brain in a homogeneous grid of 1 cm. Next, both the template and the grid were linearly transformed to fit the head shape of each subject and a local spheres approach was used to define the leadfields and fit the headshape of each subject in the vicinity of each sensor. Using the computed leadfield and the average of all the covariance matrices for each segment, we computed the spatial filter coefficients, which were used to estimate the source-space time-series for each source in all segments. 1210 source positions belonging to 80 regions of interest defined by the automated anatomical labeling atlas^38^ were identified. The remaining sources are not part of cortical regions defined by the atlas (i.e., white matter, CSF, or subcortical regions) and thus are not considered as source generators of MEG signal^39^.

To determine the individual alpha peak (IAF) we used a fast-processing algorithm immediately after the acquisition of the *pre1* and *Pre2* recordings. Some steps of the aforementioned preprocessing pipeline were omitted to obtain the IAF for the following stimulation in said frequency. Specifically, no tSSS or movement compensation method was applied, and manual revision of artifacts was skipped as well. ICA was used to remove physiological artifacts and contaminated segments were manually removed. The power spectrum for each magnetometer was calculated using DPSS, and then averaged. The resulting spectra were visually inspected, and the frequency of the power peak in the alpha band (8–12 Hz) was defined as the IAF. The IAFs from sessions *Pre1* and *Pre2* were averaged to determine the final frequency for the neurostimulation procedure.

### Neurostimulation

The neurostimulation sessions utilized bipolar tACS with two conductive rubber electrodes (7 × 5 cm) positioned at Cz (midline central) and Oz (midline occipital). A microprocessor-controlled device, the NeuroConn DC-StimulatorPlus (Neurocare, Ilmenau, Germany), was used for delivering alternating current. The electrodes were covered with sponges soaked in saline solution. For the *stim* group, stimulation at the IAF was applied during a 20-minute session with a current intensity of 3 mA peak-to-peak. In contrast, participants in the *sham* group were only exposed to stimulation during the fade-in and fade-out periods, each lasting 30 seconds. Figure 1 shows the maximum elicited electric field as simulated with ROAST^40^ and an example of the setting in a participant. After completing the stimulation and recording protocols, participants responded to a questionnaire on potential adverse effects of tACS^41^.

**Figure 1.**
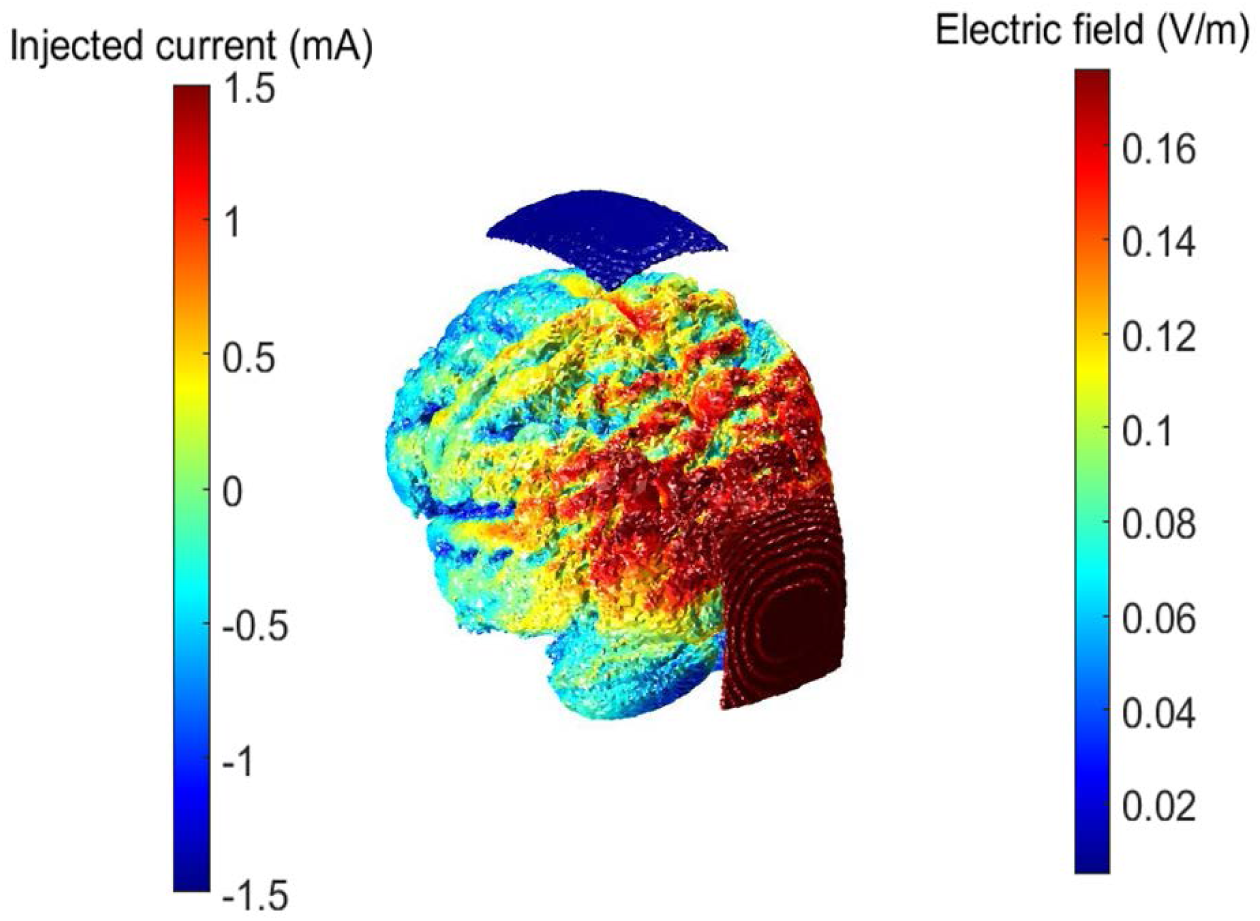
tACS maximal electric field distribution and example of the setting in a patient. Electric field distribution was estimated with ROAST.

### Functional connectivity

FC was estimated through phase and amplitude synchrony parameters after zero-phase filtering the data in the IAF ± 2 Hz frequency band through a finite impulse response filter with an order of 1800, designed with a Hamming window.

For phase-derived connectivity, the corrected imaginary phase locking value (ci-PLV^42^) was used because of its insensitivity to volume conduction, PLV’s test-retest reliability^35^ and ci-PLV’s comparability to other phase-lag connectivity measures at group-level repeatability^43^. On the other hand, amplitude-derived connectivity was calculated through the amplitude envelope correlation with leakage correction (AEC-c^44^) between the time series of each pair of the 1210 cortical sources. After orthogonalization, making it insensitive to volume conduction, the absolute value of the Pearson correlation between the time series’ envelopes was used. AEC-c was chosen because of its reliability, and within and between subject consistency^43^. Both of these FC parameters range from 0 to 1, that is, no connectivity to full connectivity.

We calculated AEC-c and ci-PLV in each of the defined signal segments, and then averaged over those, obtaining the so-called static FC (Figure 1A), resulting in connectivity matrices of dimension 1210 sources × 1210 sources. These matrices were then reduced to 40 ROIs × 40 ROIs by grouping sources according to their position in the automated anatomical labeling atlas^38^, resulting in 1210 cortical sources distributed across 40 regions of interest (80 unilateral regions grouped in bilateral analogues).

### Statistical analyses

Initially, demographic characteristics were evaluated, using an independent sample t-test to compare age and IAF, as well as a chi-square test to compare sex proportion between each of the groups. We also performed a baseline (Pre2) comparison of FC at the PCU and the DMN between the *stim* and *sham* groups.

To evaluate the influence of the stimulation protocol on FC, we performed two types of t-tests: 1) independent samples t-tests to compare the FC changes in *stim* versus *sham* groups, using a measure of their FC ratio of change; 2) repeated measures t-test to evaluate the change in FC between the Pre2 and Post per experimental group.

For the former, FC ratio of change was calculated by dividing the FC of the *Post* session of each subject by that of the *Pre2* session. Next, we calculated the logarithm of the ratio to eliminate the statistical asymmetry of ratios and obtain statistical distributions centered in 0, where negative values represent decreases and positive values represent increases over recording sessions:

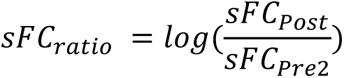

As a clarification, all analyses were performed comparing *stim* vs *sham* groups, and *Post* vs *Pre2* stages, so that a positive Cohen’s d or t-statistic would represent increased FC values in the *stim* group or *Post* stage.

These tests were applied to different sets of FC depending on the region or network to be analyzed. We evaluated the PCU and the DMN as sources, and the PCU, DMN and whole brain (i.e., AAL) as connection targets giving rise to the comparisons listed in Table 1. The DMN was defined as including the following AAL regions^45^: superior frontal gyrus (SFG), gyrus rectus (Rectus), hippocampus (Hip), anterior cingulate cortex (ACC), posterior cingulate cortex (PCC), inferior parietal gyrus (IPG), and the PCU. Each set of tests, corresponding to the area investigated (PCU or DMN), were FDR corrected with a q = 0.1 to compensate for the multiple comparisons.

**Table 1.** Set of initial statistical comparisons for the assessment of FC reduction due to tACS. This table summarizes the tests carried out initially in this study. Pre-Post cells indicate repeated measures t-test to assess FC changes in either *Stim* or *Sham* groups, and Stim-Sham cells indicate ratio t-test to evaluate differences in FC change between the *Stim* and *Sham* groups.

|  |  | To |  |  |  |  |  |
| --- | --- | --- | --- | --- | --- | --- | --- |
|  |  | PCU |  | DMN |  | AAL |  |
|  |  | Stim | Sham | Stim | Sham | Stim | Sham |
| From | PCU | Pre-Post | Pre-Post | Pre-Post | Pre-Post | Pre-Post | Pre-Post |
|  |  | Stim-Sham |  | Stim-Sham |  | Stim-Sham |  |
|  | DMN |  |  | Pre-Post | Pre-Post | Pre-Post | Pre-Post |
|  |  |  |  | Stim-Sham |  | Stim-Sham |  |

In order to evaluate which areas of the DMN drive the differences in connectivity change between the *stim* and *sham* groups, we conducted post-hoc t-tests comparing the FC ratio at each link between any two areas belonging to the DMN. Given the high number of comparisons, post-hoc results were FDR corrected with a q of 0.1 as well.

Lastly, we aimed to study the relationship between the FC ratios and simultaneous alterations in relative power given the previously suggested capacity of tACS to alter power^18^. To do this, we performed a Pearson’s correlation between the FC ratio of those DMN links that showed significant differences in the *stim-sham* comparison after FDR correction, and the ratio of IAF ± 2 relative power of the regions that make up the link (e.g. supposing the ACC-PCC link’s ratio showed significant differences between groups, the FC ratio would be correlated with the power ratio of both the ACC and PCC), independently. This analysis was carried out separately for the *stim* and *sham* groups, and their rhos were compared through a z-test on Fisher z-transformed correlation coefficients^46^.

### Simulation of tACS phase-shifts effects on FC

We performed an ad-hoc theoretical analysis to investigate the potential effects of cortical morphology over FC modulation. Given our current stimulation protocol and cortical folding, the neurons of certain regions may receive the stimulation in antiphase, as the electric field direction may result anti-parallel to some cortical columns (i.e., hyperpolarizing) and parallel to others^18^ (see Figure 7). To evaluate the effect of this phase disparities over FC, we simulated two MEG signals through filtered white noise and calculated their initial connectivity through PLV and AEC. Afterwards, we added a sinusoidal stimulus to each of them with different phase shifts and calculated again PLV and AEC. This procedure was repeated 100 times.

A repeated measures t-test was performed to compare the connectivity values between the cases of no-stimulation, sinusoid stimulation with 0 phase shift, and sinusoid stimulation with π phase shift. Supplementary material 1 explains the complete methodology for the simulation process.

## Results

### Sample demographics and adverse tACS effects

Sample demographics and variables of interest used in this study are shown in Table 2, and reported tACS effects are depicted in Supplementary Table S1. No statistically significant differences were observed between the groups either in demographics or tACS adverse aftereffects, or in baseline FC.

**Table 2.** Sample demographics. The differences in each of the demographic variables are shown in the two rightmost columns, displaying the t-statistic or chi-square statistic for continuous and categorical variables, respectively, and p-value. IAF: Individual Alpha Frequency.

|  | *Stim* | *Sham* | Stat | p-value |
| --- | --- | --- | --- | --- |
| Age | 33.4 ± 8.4 | 32.2 ± 9.0 | 0.3940 | 0.7302 |
| Sex | 9M / 5F | 6M / 6F | 0.5403 | 0.4623 |
| IAF | 10.3 ± 0.9 | 10.5 ± 1.3 | -0.5741 | 0.5713 |
| Pre2 PCU<br>AEC-c | 0.210 ±<br>0.007 | 0.0210 ±<br>0.007 | 0.0377 | 0.9702 |
| Pre2 DMN<br>AEC-c | 0.202 ±<br>0.004 | 0.0203 ±<br>0.003 | -0.3886 | 0.7010 |
| Pre2 PCU<br>ci-PLV | 0.176 ±<br>0.008 | 0.182 ±<br>0.008 | -1.9242 | 0.0663 |
| Pre2 DMN<br>ci-PLV | 0.169 ±<br>0.007 | 0.170 ±<br>0.005 | -0.2887 | 0.7753 |

### AEC-c at the PCU is reduced by tACS

To assess the effect of tACS on the PCU’s FC, we performed independent samples ratio t-tests comparing the FC change between *stim* and *sham* groups. Then, we performed repeated measures t-tests to evaluate statistically the change from *Pre2* to *Post* stimulation for each group. The FC values were evaluating three relationships: PCU-PCU, PCU-DMN, and PCU-AAL. FDR correction (q = 0.1, *n_comparisons_* = 18) yielded a critical p-value of 0.0425. Figure 2 shows the results for these analyses.

**Figure 2.**
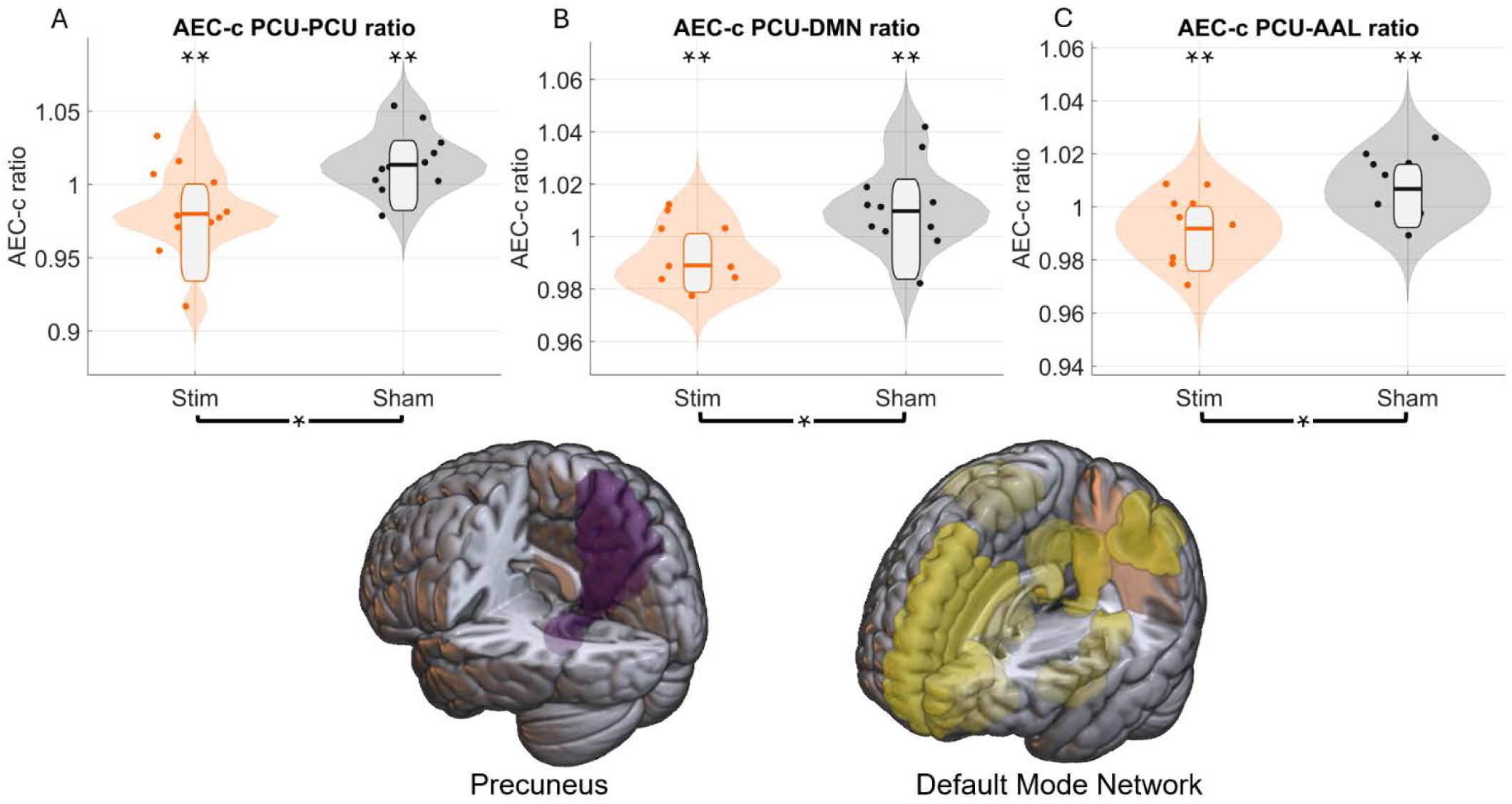
tACS reduces PCU’s connectivity. Violin plots show the *Post/Pre2* FC ratio for PCU intraconnectivity (A), between the PCU and the DMN (B), and between the PCU and the rest of the brain (C), with orange violin plots corresponding to the *stim* group and the grey ones to the *sham* group. Single asterisks (*) and brackets below the graphs indicate significant between-group differences in the FC ratio, while double asterisks (**) above each violin plot indicate significant intra-group pre/post differences. Brain images at the bottom of the figure represent the areas of the PCU and the DMN included in the study. Importantly, the PCU was not included in the DMN for this analysis.

The independent samples t-tests found significant decreases of AEC-c for all spatial relationships (Figure 2): PCU-PCU [t-stat = -3.3726, d-Cohen = -1.3268, p-value = 0.0025], PCU-DMN [t-stat = -3.6406, d-Cohen = -1.4322, p-value = 0.0013], and PCU-AAL [t-stat = -3.6165, d-Cohen = -1.4227, p-value = 0.0014]. Repeated-measures t-test showed significant decreases *Pre2*-*Post* of AEC-c for the *stim* group in every spatial relationship (Figure 2): PCU-PCU [t = -2.3152, d-Cohen = - 0.3530, p-value = 0.0376], PCU-DMN [t = -2.7491, d-Cohen = -0.3374, p-value = 0.0166], and PCU-AAL [t = -2.8251, d-Cohen = -0.3117, p-value = 0.0143]; while significant increases were observed for the *sham* group: PCU-PCU [t-stat = 2.5845, d-Cohen = 0.2575, p-value = 0.0254], PCU-DMN [t-stat = 2.4821, d-Cohen = 0.3231, p-value = 0.0305], and PCU-AAL [t-stat = 2.2942, d-Cohen = 0.2266, p-value = 0.0425].

No significant effects of tACS on the PCU FC were found through ci-PLV. Disclosure of all statistical results can be found in supplementary table S2.

### DMN’s functional connectivity decreases through tACS

Now, we applied independent samples t-tests to assess the differences in the ratio of change between the groups. Then, repeated-measures t-tests were performed to evaluate Pre2-Post differences in the FC of the DMN in each group. Two spatial relationships were considered, including the DMN-DMN and the DMN-AAL. FDR correction (q = 0.1, *n_comparisons_* = 12) yielded a critical p-value of 0.0574. Given that FDR’s critical-value is higher than 0.05, only results under a p-value of 0.05 will be considered significant.

The comparisons of the ratio of change in AEC-c between experimental conditions showed a significant difference between both groups in the two spatial relationships (Figure 3): DMN-DMN [t-stat = -2.8614, d-Cohen = -1.1257, p-value = 0.0086], and DMN-AAL [t-stat = -2.7104, d-Cohen = -1.0663, p-value = 0.0122]. In consonance with the previous results for the PCU, repeated-measures t-tests revealed significant decreases in AEC-c between the *Post* and *Pre2* sessions in the *stim* group for each of the two spatial relationships (Figure 3): DMN-DMN [t-stat = -3.9109, d-Cohen = -0.6383, p-value = 0.0018], and DMN-AAL [t-stat = -3.2451, d-Cohen = - 0.3966, p-value = 0.0064]. However, no significant differences were observed in the *sham group:* DMN-DMN [t-stat = 0.6828, d-Cohen = 0.1456, p-value = 0.5089]; DMN-AAL [t-stat = 1.1133, d-Cohen = 0.2365, p-value = 0.2893].

**Figure 3.**
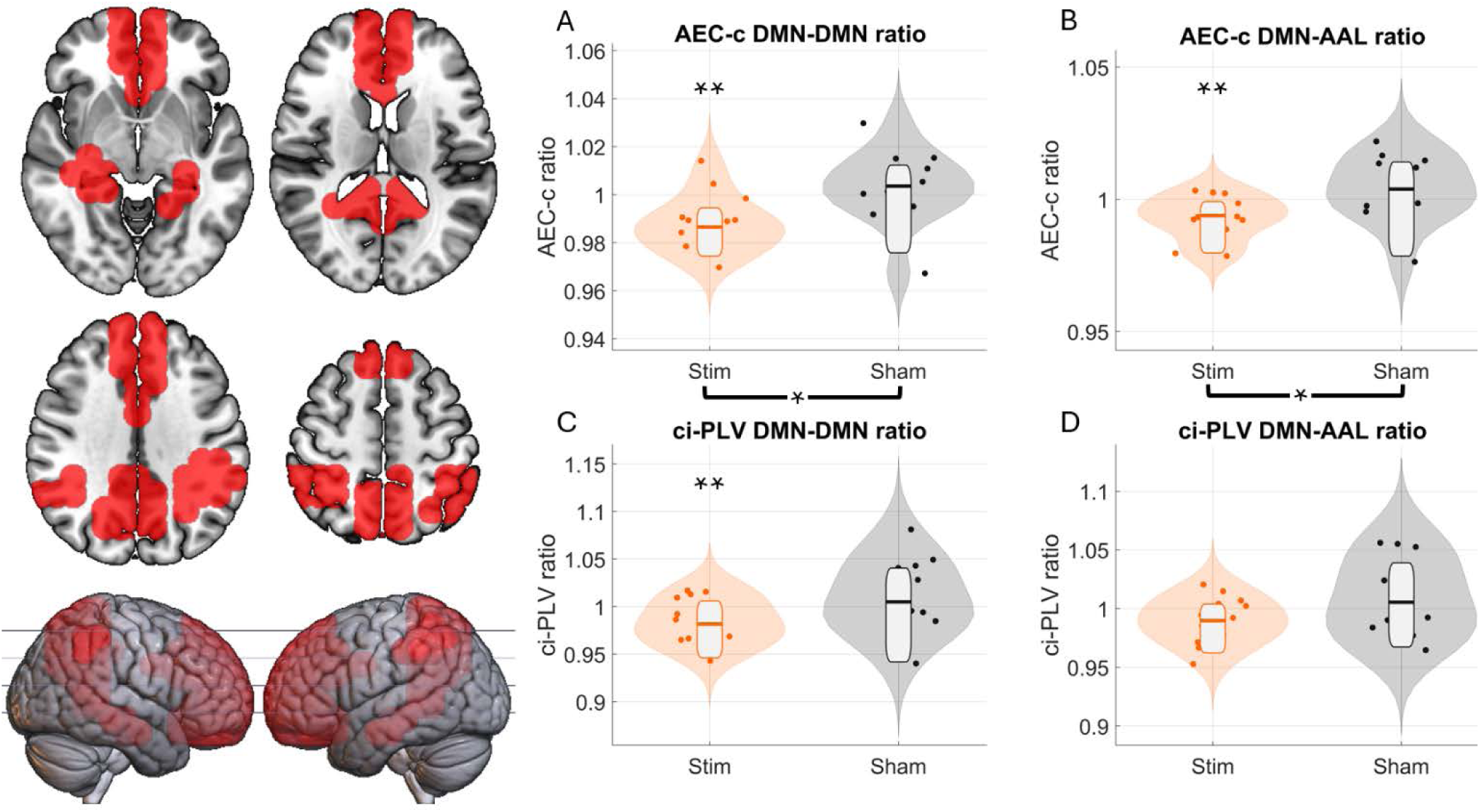
DMN connectivity decreases due to tACS. Violin plots show the *Post/Pre2* FC ratio of DMN intraconnectivity (A and C), and between DMN and the rest of the brain (B and D). Significant reductions of FC between the *stim* and sham groups were detected with AEC-c (A and B), as noted by the single asterisks (*) and the brackets below the graphs. At the same time, the *stim* group showed significant intra-group decreases of AEC-c and ci-PLV for DMN intraconnectivity (A and C, respectively), and of AEC-c between the DMN and the rest of the brain (B), as indicated by the double asterisks (**) above their violin plots.

On the other hand, changes in the ci-PLV of the *stim* group, relative to those in the *sham* group, showed decreases in the former that were close to statistical significance, both in DMN-DMN [t-stat = -2.0171, d-Cohen = -0.7935, p-value = 0.0550] and DMN-AAL [t-stat = -1.8447, d-Cohen = -0.7257, p-value = 0.0775], as depicted in Figure 3C and 3D. Repeated-measures t-tests revealed significant ci-PLV decreases between the Post and *Pre2* sessions in the *stim* group for DMN-DMN [t-stat = -2.6658, d-Cohen = -0.4207, p-value = 0.0194], and close to significance In the case of the DMN-AAL connectivity [t-stat = -2.0841, d-Cohen = -0.2776, p-value = 0.0574]. No significant differences were observed in the *sham group* (DMN-DMN [t-stat = 0.8127, d-Cohen = 0.2132, p-value = 0.4336]; DMN-AAL [t-stat = 0.9598, d-Cohen = 0.2091, p-value = 0.3578]).

### tACS prominently decreases posterior-posterior and antero-posterior hub links of the DMN

Additional statistical tests were performed to identify the links of the DMN driving the results presented in the previous sections. Given the high number of comparisons (*n_comparisons_* = 28), results were FDR-corrected with a q = 0.1.

Out of the 28 possible links, 7 showed statistically significant differences using AEC-c including Hip-ACC, IPG-IPG, PCC-PCC, PCU-Rectus, PCU-PCC, PCU-ACC, and PCU-PCU (Figure 4). These differences were in the same direction as reported previously (i.e., a decrease in the FC of the *stim* group). No ci-PLV link difference was significantly different after FDR. Table S3 and Table S4 include a complete report for the 28 possible links in both connectivity metrics for AEC-c and ci-PLV, respectively.

**Figure 4.**
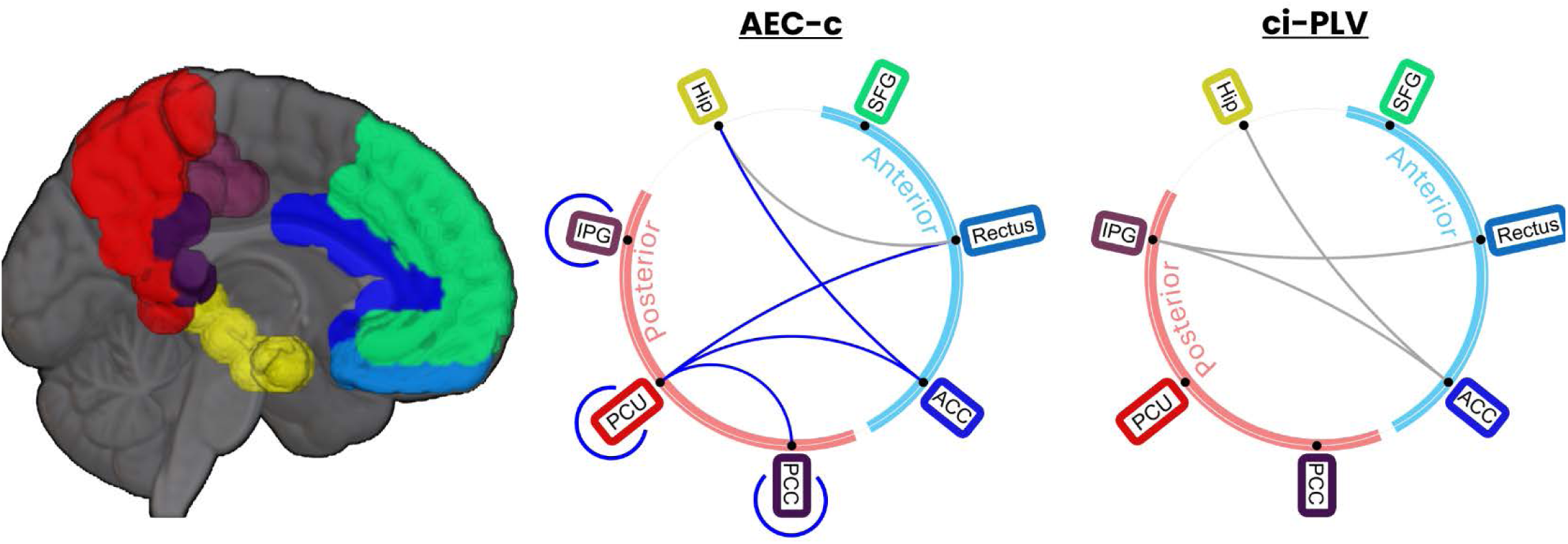
DMN links show a significant decrease in connectivity after tACS. Ratio t-tests revealed 8 DMN links showing a significant AEC-c reduction and 3 showing a significant ci-PLV reduction when comparing *stim* and *sham* groups. Of those, 7 AEC-c links still showed significance after FDR correction (q = 0.1), while no ci-PLV links survived. Links significant after FDR correction are indicated in blue, while those that lost significance are indicated in grey.

### Functional connectivity of posterior areas correlates with power change

To understand the relationship between FC and power changes elicited by the stimulation, we performed a post-hoc analysis correlating the AEC-c value - of the significant links reported in the previous section - and the relative power of the IAF ± 2 Hz of each area involved. This represented 11 Pearson’s correlations per group, as self-connections (e.g., PCU-PCU) involved one test, while inter-regional connections (e.g., PCU-PCC) involved two.

Out of the seven links studied, two of them showed significant negative correlations between FC and normalized power in the *stim* group: PCU-PCU and PCU-PCC correlating with both regions relative power (Figure 5). Nevertheless, the only case that exhibited a significantly different correlation pattern between *stim* and *sham* groups was that between the PCC-PCU’s FC and the normalized power of the PCU [z-stat = -2.3703, p-value = 0.0178]. Importantly, no differences in the relative power in the IAF ± 2 Hz band were found between the *stim* and *sham* groups in any of the studied regions.

**Figure 5.**
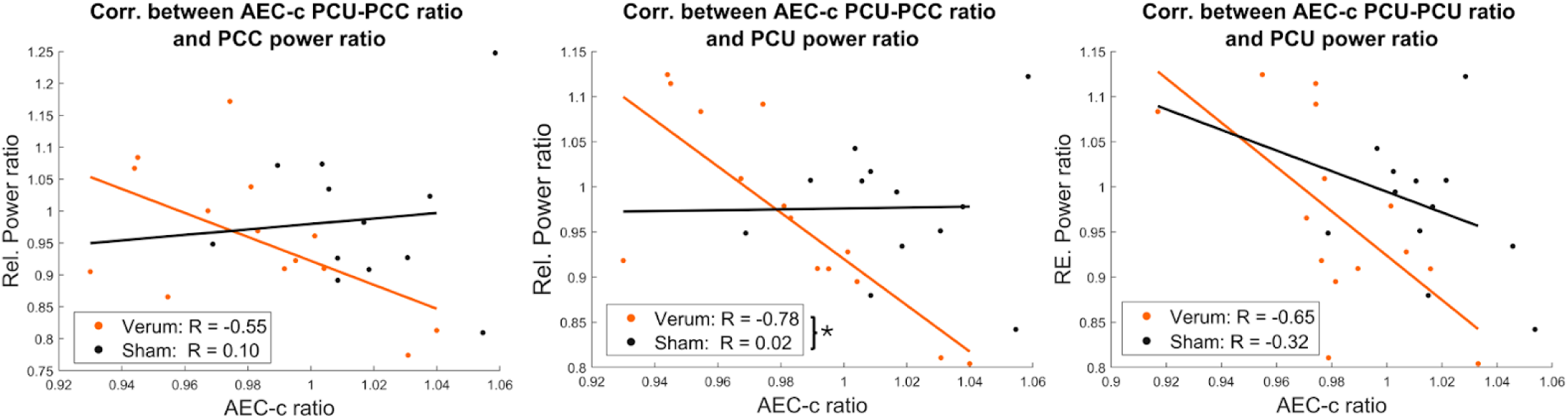
Negative correlations between AEC-c FC ratio in posterior areas and IAF ± 2 Hz relative power. Significant correlations between AEC-c and relative power in the *stim* group (orange) were found in posterior regions, while these correlations were nonsignificant in the *sham* group (black). However, only the correlation between the PCC-PCU and the IAF ± 2 Hz relative power at the PCU was able to significantly differentiate between the *stim* and *sham* groups, as indicated by a single asterisk (*).

### Simulating tACS-induced phase disparities explain the reductions of FC

As stated in the introduction, while we expected to observe an increase in FC after the stimulation protocol, the results showed a generalized decrease in FC in the *stim* group. In order to understand further these results, we studied the potential interaction of cortical morphology with the stimulation protocol. Given the brain cortical folding, the neurons of certain regions may receive the stimulation in anti-phase, as the electric field direction may result antiparallel to some cortical columns (i.e., hyper-polarizing effect) and parallel to others^18^ (see Figure 7). Therefore, we performed a theoretical experiment by introducing phase disparities in two simulated MEG signals and measuring the resulting FC.

We found that any phase shift between the stimuli deviating from 0 or 2π phase produced a decrease in connectivity both for PLV and AEC. Specifically, significant differences in connectivity values both for PLV [t-stat = 4.7082, p-value = 0.0011], and AEC [t-stat = 7.3485, p-value < 0.0001] emerged when comparing the cases of no stimulation and anti-phase stimulation (phase = π). No significant differences were found when comparing connectivity in the case of no stimulation and stimulation with in-phase sinusoids both for PLV [t-stat = -1.4465, p-value = 0.1819] and AEC [t-stat = -1.9688, p-value = 0.0805].

## Discussion

In this study, we assessed the influence of a parieto-occipital IAF-tACS protocol on FC measured by MEG. Both ci-PLV and AEC-c were calculated and then compared through the ratio between pre- and post-stimulation sessions for the *stim* and *sham* groups. The FC ratio for the AEC-c was significantly lower in the *stim* group than in the *sham* group, both in the PCU and in the whole DMN, and in all cases the AEC-c in the *stim* group decreased significantly between pre- and post-sessions. Furthermore, we studied the links of the DMN that changed with the stimulation, finding a prominent decrease in AEC-c over posterior areas and in the connectivity between postero-anterior hubs of the DMN. Finally, we found negative correlations relating the change in AEC-c in posterior DMN areas and the change in power in those same areas. These results complement and expand previous literature by assessing the influence of personalized tACS on FC in healthy participants.

These results do not support our hypothesis of enhanced FC after tACS, which was based on previous literature^23,24,47^. On the one hand, we hypothesize that the reductions in FC found in this study might be explained by the interaction between the electric field direction and the position of pyramidal neurons along the folded cortex. Where the electric field direction is aligned parallel to the body axis of pyramidal cells, depolarization will be elicited. Inversely, where the alignment is antiparallel, hyperpolarisation occurs. This introduces phase disparities in the stimulation induced and its intensity (Figure 7).

To illustrate the possible effects of these phase disparities over both phase and amplitude FC, we performed an ad-hoc analysis simulating the influence of tACS over physiological MEG signals. We found that any phase shift between the stimulatory sinusoids deviating from 0 or 2π phase produced a decrease in connectivity both for PLV and AEC (Figure 6). For phase-based connectivity, the phase shift between the stimulation signals changes the spectral contribution of the stimulation frequency to the original signals, depending on the constructive or destructive interaction with the physiological signals’ activity at the stimulation frequency. The differences in spectral density will result in different instantaneous frequency probability distributions for each of the signals, making it less likely for them to maintain a constant phase difference, and thus decreasing the PLV. On the other hand, the phase difference between the stimulatory sinusoid waves modifies the envelopes differently, reaching a maximum misalignment of the original physiologic signals’ envelopes when the phase shift is that of π, consequently reducing their AEC. The interpretation of these simulation results should be considered with caution as many assumptions have been made. For instance, we have assumed a straightforward behaviour and influence of tACS as sinusoidal signals over neuronal physiological signals, and considered white noise as a plausible proxy for neuronal activity. This theoretical experiment adds to the understanding of the results obtained in this study.

**Figure 6.**
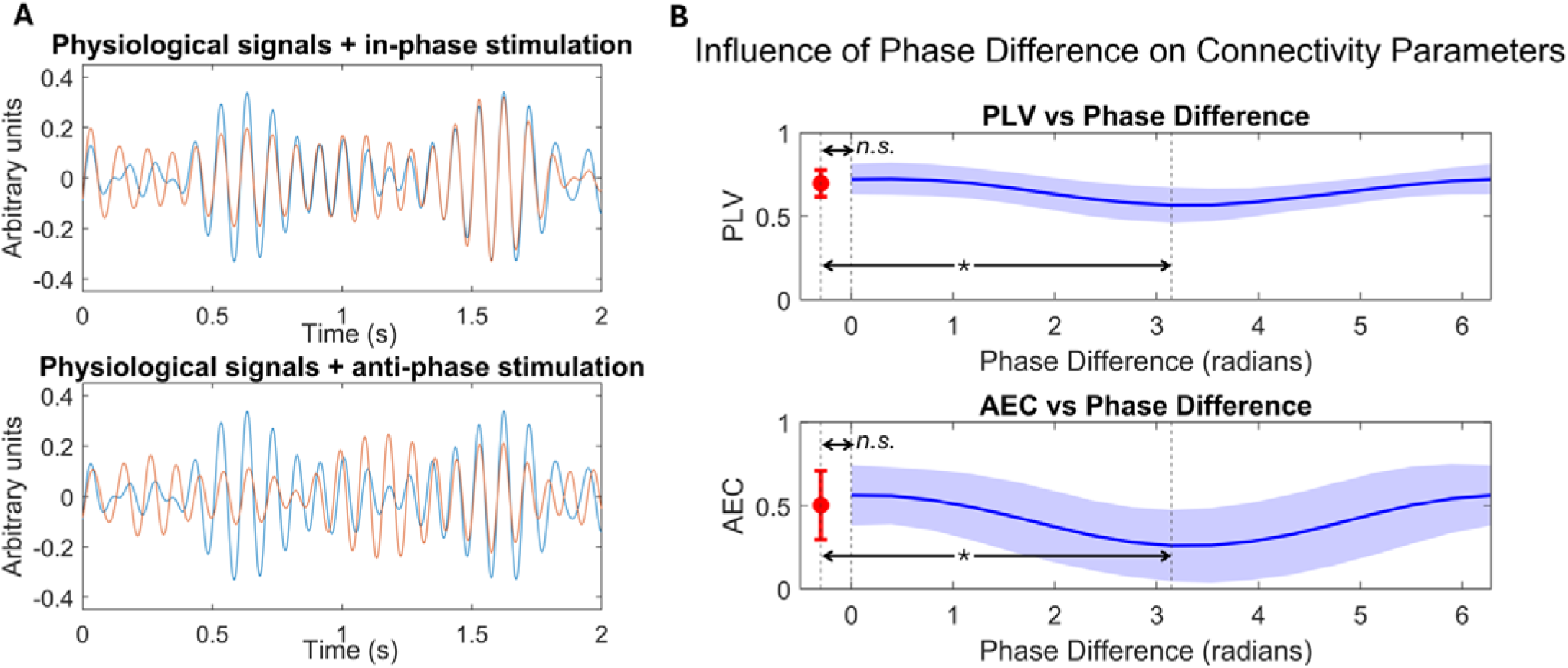
Results of the simulation of the effect of phase-shift tACS influence on FC. A) Physiological signals being stimulated with in-phase stimulation (top), and with antiphase stimulation (bottom). B) Relation between PLV and AEC and the phase difference between stimulation sinusoids. Significant reductions between antiphase stimulation and initial connectivity were found and are indicated by an asterisk. N.s: non-significant.

**Figure 7.**
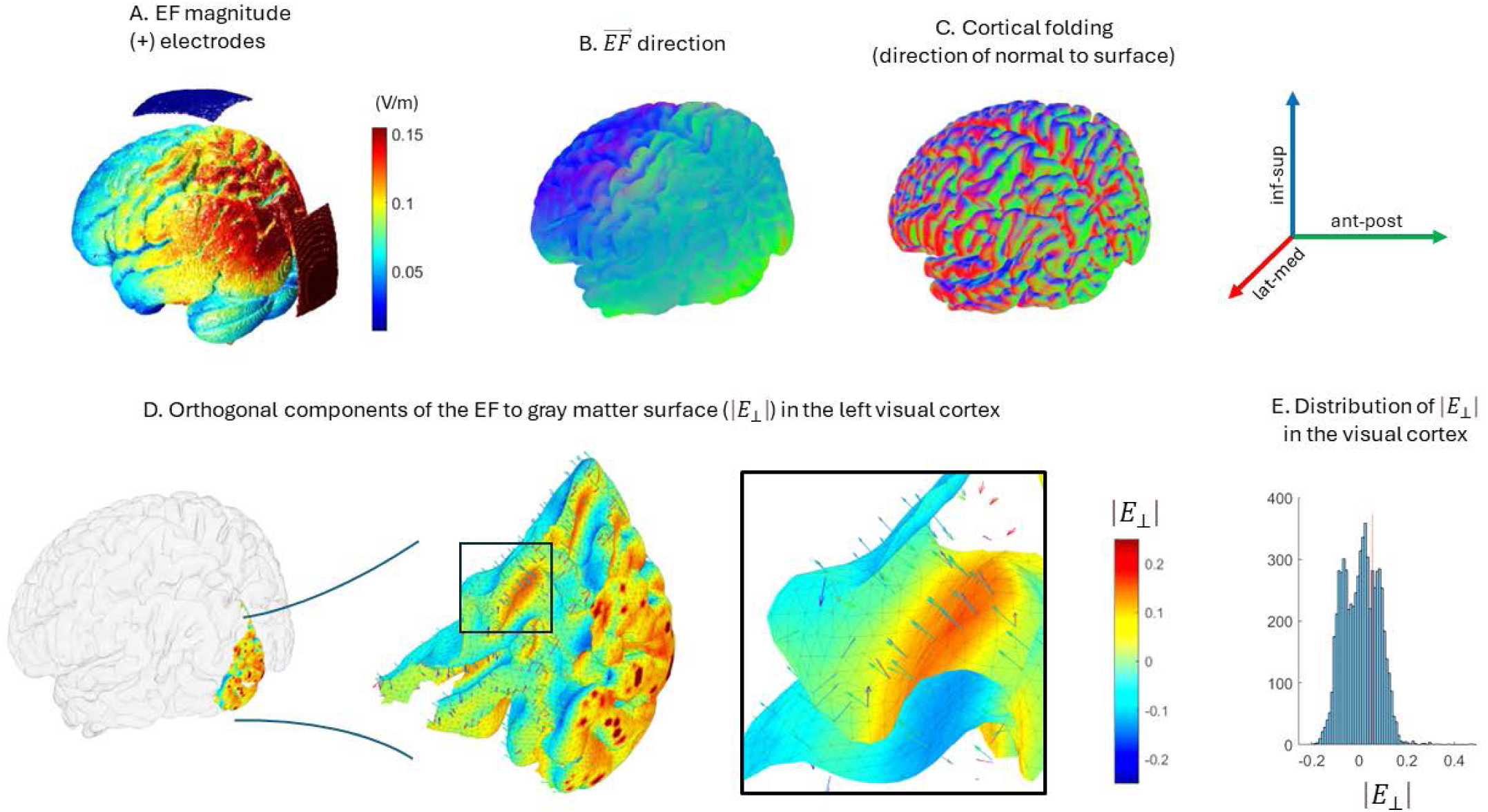
Orthogonal components of the electric field with respect to the white-matter surface. A) Electrode location and electric field magnitude over the brain. B and C show the direction of both the electric field given the protocol (B) and the direction of the normal component to white matter surface in the folded cortex (C). D) Distribution of orthogonal components of the electric field in the visual cortex. A sample of vectors is shown indicating both the direction of the electric field and the normal component to the surface. The surface colour indicates the magnitude of the orthogonal component (red-depolarizing, blue-hyperpolarizing). E) Distribution of magnitudes of the orthogonal components of the electric field for the visual cortex. Note that in the same region coexist positive and negative values, and therefore, anti-phase stimulation effects.

An equally plausible explanation for our divergent results when compared to previous findings by others could stem from the lack of comparability in the field^48^, due to the use of different stimulation protocols, spatial targets, or means to measure the stimulation effects. The study performed by Schwab et al.^23^ reported an increase in alpha functional connectivity for occipitoparietal in-phase tACS when compared to anti-phase and jittered-phase tACS, but a sham group was not included, and thus our results are not comparable with theirs. Additionally, their tACS protocol was administered through HD-tACS, where the stimulation electrodes are surrounded by returning electrodes in an effort to focalize the elicited electric field. This is especially relevant, as the phase differences induced in the neural populations in the cortex would be highly different between protocols. This is the same case as in Clancy et al.^24^, which applied HD-tACS to occipito-parietal areas using the IAF as stimulation frequency, and additionally used Granger causality, a different FC parameter than ours, to measure changes in connectivity. In the work by of Schouwenburg et al.^47^, both increases and decreases in FC were observed in different areas of the brain, and a second study of the same group could not replicate the results found in the first one^49^. Finally, other studies based on fMRI-FC show inconclusive results, with some finding increases in FC after tACS stimulation^50,51^ and other finding FC reductions^25,52^. Consequently, there is a need for replication studies and a standardization of the stimulation protocols to evaluate the effects of tACS on FC.

For a better understanding of the electrophysiological phenomena underlying the use of tACS, we conducted an additional study on the joint behaviour of FC and power in the DMN links with significant changes after tACS. This revealed a consistent pattern of negative correlations between the change of AEC-c in the PCU-PCC link, and power of these two regions for the *stim* group. Only the correlation with the PCU’s power distinguished the behaviour between the *sham* and *stim* groups, possibly due to the reduced statistical power of the sample. Aligned with the previous explanation for the FC reductions, tACS could introduce non-uniform phase distributions in the cortex, favouring a decoupling of phases and envelopes, but still entrain the activity of the neuronal populations in the same stimulation frequency. This would consequently increase the activity within the frequency range, increasing power^18,20^ but reducing FC as shown previously.

Interestingly, the reduction in PCU-ACC and PCU-PCC FC is of special interest in the context of Alzheimer’s Disease (AD). Specifically, an early rise in FC between these areas is found in individuals at risk of disease development due to their family history^53,54^. Moreover, the whole DMN is affected in the disease as its function, metabolic activity and structure become altered through the course of the disease^55^. Some of these regions, such as the PCU, behave as hubs of the network and become vulnerable to the progression of the disease^56^. Particularly, these regions show increased excitability and connectivity in the earliest stages of Aβ accumulation, which intensifies the severity of the disease^57–59^. Therefore, reducing or limiting the initial increase in neuronal excitability and connectivity through a non-invasive technique such as tACS could pose as a potential preventive treatment in prodromal stages of AD. Further investigation in this direction is needed, as well as in the cognitive effects of the proposed tACS protocol.

Some limitations of the present experiment must be noted. Firstly, while comparable to previous studies in the field, the sample size of our experiment was modest, with a total of 26 participants. This limited the possibility of introducing potential confounding variables as covariates. Nevertheless, no significant differences in any of the demographic variables between the *sham* and *stim* groups were observed. Future studies should consider larger sample sizes to increase statistical power. Importantly, even though physiological relative IAF ± 2 Hz power was not used as a covariate, since there were no significant differences in relative power between groups, the possibility for power to constitute a confounding factor in our study is slim. Secondly, the temporal length of the protocol could have induced fatigue in the participants, influencing the results in terms of power and FC^60^. Finally, it is important to note that the stimulation setup with squared patches is limited in its spatial focality, inducing current into large areas of the brain.

Our investigation lacks behavioral evaluation and consequently did not address the cognitive relevance of the findings. Also, the current literature presents elevated variability in the methods and stimulation used, yielding non-comparable results between studies. Therefore, future research should focus on standardizing methodologies, replicating previous results, including neuropsychological assessments, combining electrophysiological and fMRI measurements, as well as using personalized anatomical targeting to reduce inter-subject variability^61,62^. The use of computational modeling to systematically simulate and study the differences between stimulation protocols would also contribute to knowledge convergence and experiment reproducibility in the field.

## Conclusion

To conclude, this is, to the best of our knowledge, the first study assessing both offline phase- and amplitude-based FC changes induced by personalized IAF-tACS with MEG in healthy participants. Our results suggest that tACS reduced AEC-c and, to a lesser extent, ci-PLV of both the PCU and the DMN through IAF-tACS over occipito-parietal areas. We hypothesize that this effect is mediated by the alternating in-phase and anti-phase neuromodulation that tACS produces in opposite oriented cortices, which might lead to the reduction of both phase and amplitude synchrony. Lastly, we find that the changes in FC between posterior areas negatively correlated with power increases in the same frequency band. Our findings highlight a potential treatment for functional connectivity reduction, potentially beneficial for AD, and expand the knowledge of the possible influence of brain folding on the individual effects of tACS.

## Acknowledgements

We would like to thank all participants that were included in this study for their selfless contribution to science, making this work possible.

## Authorship statement

Fernando Maestú, Carlos J Gómez-Ariza, Martín Carrasco-Gómez, and Alberto del Cerro-León designed and conducted the study, including participant recruitment, data collection, and data preprocessing. Martín Carrasco-Gómez performed the data analysis and statistical assessment of the present study and prepared the manuscript draft with important intellectual input from Alejandra García-Colomo, Jesús Cabrera-Álvarez and Alberto del Cerro León. Carlos J. Gómez-Ariza, Andrés Santos and Fernando Maestú reviewed and aided in the final preparation of this manuscript.

## Abbreviations

FC: functional connectivity
PCU: precuneus
DMN: default mode network
IAF: individual alpha frequency
MEG: magnetoencephalography
PCC: posterior cingulate cortex
ci-PLV: corrected imaginary phase-locking value
AEC-c: leakage corrected amplitude envelope correlation
IPG: inferior parietal gyrus
ACC: anterior cingulate cortex
Hip: Hippocampus
SFG: superior frontal gyrus
Rectus: gyrus rectus

