## Supplementary material for "Individual alpha frequency tACS reduces functional connectivity across the default mode network"

### Simulation on tACS phase shift effects on functional connectivity (FC)

To study the possible effects of phase disparities introduced by tACS in the brain cortex over both phase and amplitude FC, we simulated MEG signals, adding to them two stimulatory sinusoid time series with different phase shifts, and finally calculated the phase locking-value (PLV) and the amplitude envelope correlation (AEC).

The complete process took the following steps:

1. **Configuration for the signals simulation**

% Parameters

Fs = 1000;	% Sampling rate

T = 8;	% Duration in seconds

filter_low = 8;	% Lower bound of filter

filter_high = 12;	% Upper bound of filter

filter_order = 1800;	% Filter order

sin_freq = 10;	% Frequency of the stimulation sinusoidal signals

phases = [0, pi/8, pi/4, 3*pi/8, pi/2, 5*pi/8, 3*pi/4, 7*pi/8, pi, 9*pi/8, 5*pi/4, … 11*pi/8, 3*pi/2, 13*pi/8, 7*pi/4, 15*pi/8, 2*pi]; % Different phase differences

amp_sin = 0.5	% The amplitude of the stimulation sinusoids will
be 25% of the mean amplitude of the physiological

signal

amp_noise = 0.5	% The amplitude of the noise signals will be 50% of

that of the physiological signal

The sampling rate, duration of the signal and the filter order were selected so that they would fit the parameters of the experimental signal and processing of the study. Given that the study is focused on the alpha band, the signals were filtered between 8 and 12 Hz, and the frequency for the stimulation sinusoids was set to 10 Hz, emulating the study setting. The phase vector included 17 values in the interval [0,2π].

2. **White noise signal generation**

% Generate random signals

n = Fs * T; % Number of samples

meg1	= randn(1, n);	% Basis for the physiological signals

meg2	= randn(1, n);	% Basis for the 1st noisy signal

meg3	= randn(1, n);	% Basis for the 2nd noisy signal

We generate our signals from a normal distribution centered in 0 and with a standard deviation of 1. We generate 3 signals to later create two physiological signals by combining *meg1* and *meg2,* and *meg1* and *meg3*, respectively.

3. **Filtering of the signals**

% Filter signals between 8 and 12 Hz

d = designfilt('bandpassfir', 'FilterOrder', filter_order, 'CutoffFrequency1', … filter_low, 'CutoffFrequency2', filter_high, 'SampleRate', Fs);

filtered_signal1 = filtfilt(d, meg1);

filtered_signal2 = filtfilt(d, meg2);

filtered_signal3 = filtfilt(d, meg3);

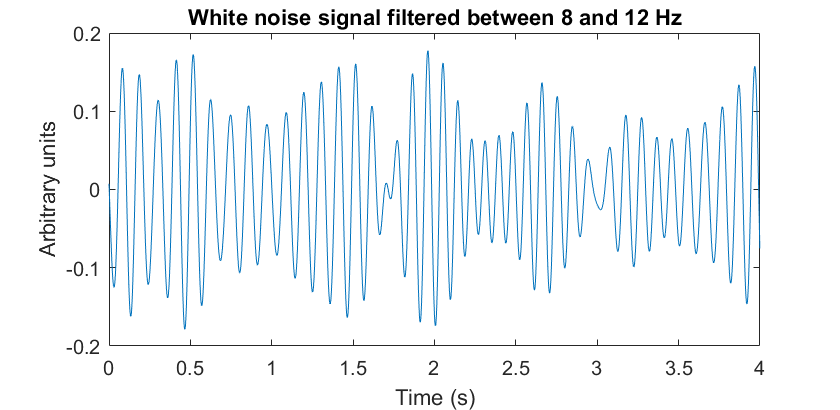

A filter is designed to have cutoff frequencies of 8 and 12 Hz, and a filter order of 1800. Then the white noise signals are zero-phased filtered using the *filtfilt* function.

4. **Padding removal**

% Remove 2 seconds of padding (2000 samples) from the beginning and end

filtered_signal1 = filtered_signal1(2001:end-2000);

filtered_signal2 = filtered_signal2(2001:end-2000);

filtered_signal3 = filtered_signal3(2001:end-2000);

2 seconds are eliminated from the beginning and the end of the signal, so there is no border effect on the simulated signals.

5. **Stimulation sinusoid amplitude determination**

amp_sine = amp_sine * mean(abs(filtered_signal1));	% 50% peak-to-peak is

25% of amplitude

The peak-to-peak amplitude for the stimulation sinusoids is 50% of the mean of the absolute value of the signal (necessary, since the estimated mean of the signal should be close to zero).

6. **Combination of white noise signals to create two physiological signals**

% Combine together original and noisy signals

combined_signal1 = filtered_signal1 + amp_noise * filtered_signal2;

combined_signal2 = filtered_signal1 + amp_noise * filtered_signal3;

Then we combine the signals, so that the resulting couple of signals have some synchronization, but they still are different. Given that amp_noise is 0.5, then the resulting signals have 66.6% in common.

7. **Time vector generation**

% Generate time vector

t = (0:length(filtered_signal1)-1) / Fs;

8. **Initialization of the PLV and AEC vectors**

% Initialize arrays to store PLV and AEC values

PLV_values = zeros(1, length(phases)+1);

AEC_values = zeros(1, length(phases)+1);

The vector where the connectivity values are will be stored have one extra space to save the FC between the signals with no stimulation involved.

9. **Initial connectivity calculation between the physiological signals**

% Calculate Phase Locking Value (PLV)

phase_diff = angle(hilbert(combined_signal1)) - angle(hilbert(combined_signal2));

PLV_values(1) = abs(mean(exp(1j * phase_diff)));

% Calculate Amplitude Envelope Correlation (AEC)

envelope1 = abs(hilbert(combined_signal1));

envelope2 = abs(hilbert(combined_signal2));

AEC_values(1) = corr(envelope1', envelope2');

The initial connectivity values are calculated from the two “physiological” signals, by calculating the average phase difference in the case of the PLV, and the correlation between the signals’ envelopes in the case of the AEC.

10. **Loop for iterative addition of differently phase-shifted stimulation sinusoids**
    a. Generation of the stimulation sinusoids

% Generate sinusoidal signals with specified phase difference

sinusoid1 = amp * sin(2 * pi * sin_freq * t);

sinusoid2 = amp * sin(2 * pi * sin_freq * t + phases(i));

b. Combination of the physiological signals and the stimulation sinusoids

% Add sinusoidal signals to filtered signals

combined_signal1 = filtered_signal1 + sinusoid1 + ampNoise * filtered_signal2;

combined_signal2 = filtered_signal1 + sinusoid2 + ampNoise * filtered_signal3;

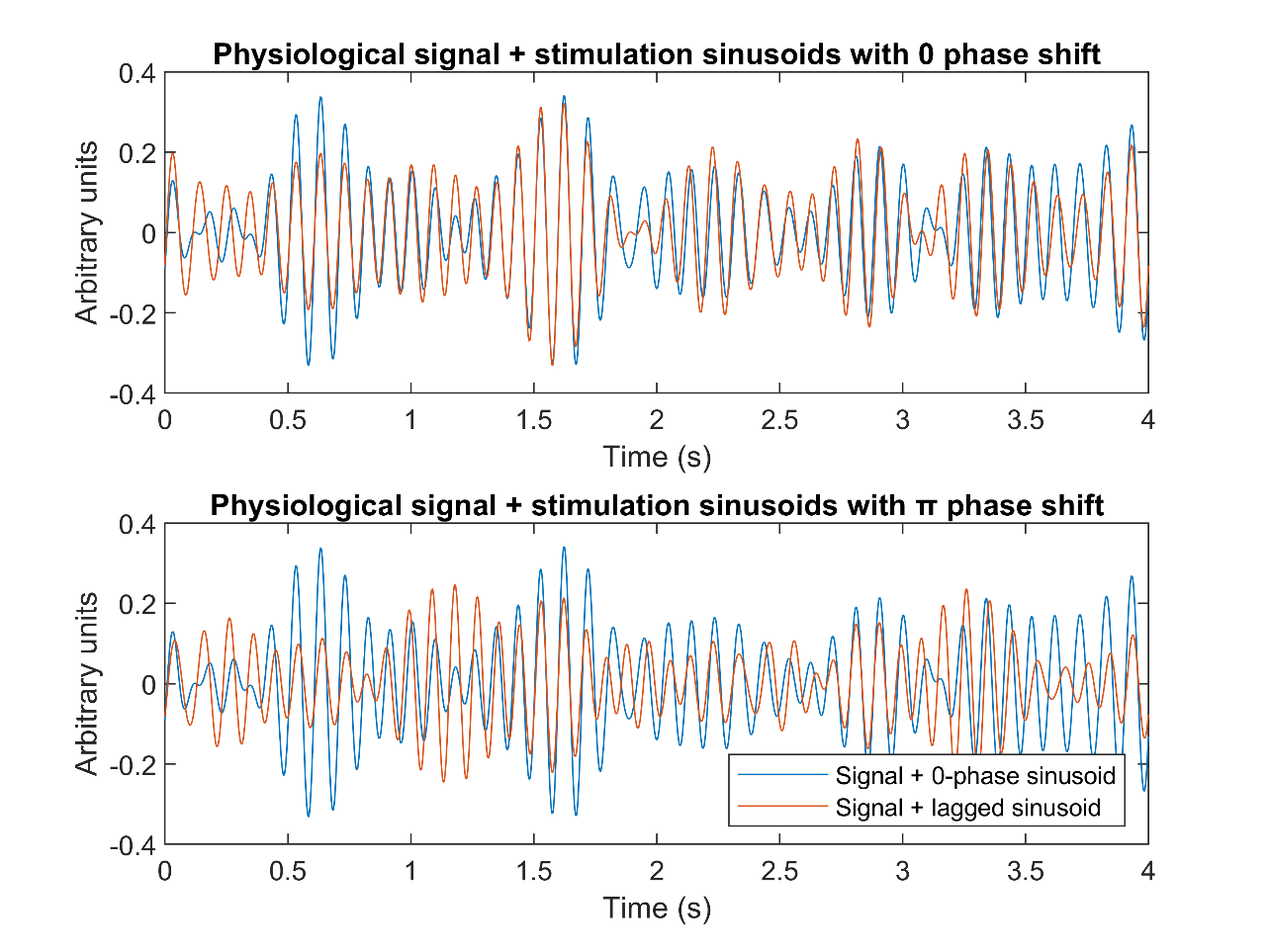

c. Calculation and storage of the PLV and AEC values

% Calculate Phase Locking Value (PLV)

phase_diff = angle(hilbert(combined_signal1)) - angle(hilbert(combined_signal2));

PLV_values(i+1) = abs(mean(exp(1j * phase_diff)));

% Calculate Amplitude Envelope Correlation (AEC)

envelope1 = abs(hilbert(combined_signal1));

envelope2 = abs(hilbert(combined_signal2));

AEC_values(i+1) = corr(envelope1', envelope2');

This process was repeated over 100 times, so that we could obtain distributions of the influence of phase shifts over FC. Finally, repeated measures t-test were carried out between the no-stimulation condition and the in-phase stimulation (phase shift between signals = 0), and between the no-stimulation condition and the anti-phase stimulation (phase shift between signals = π).

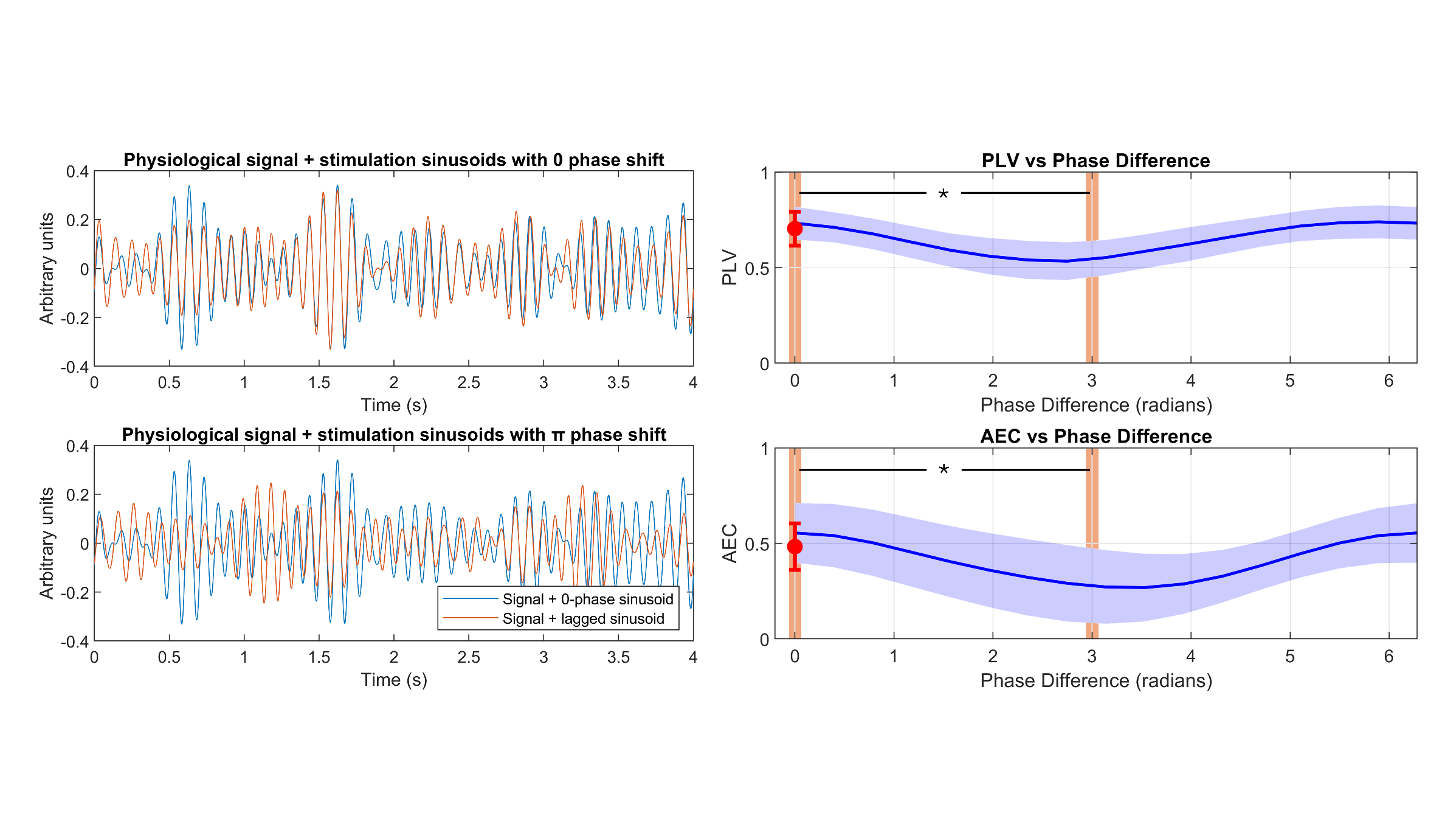

### Supplementary tables

**Table S1. Distribution of reported adverse effects of tACS by group.**

|  | Stim | Sham | t-stat | p-value |
| --- | --- | --- | --- | --- |
| Headache | 1.28 ± 0.61 | 1.17 ± 0.39 | 0.5804 | 0.567 |
| Neckache | 1.07 ± 0.27 | 1.08 ± 0.29 | -0.1091 | 0.914 |
| Pain in the scalp | 1 ± 0 | 1 ± 0 | 0 | 1 |
| Tingling | 2 ± 0.78 | 1.83 ± 0.72 | 0.5614 | 0.5797 |
| Stinging sensation | 1.64 ± 0.63 | 1.58 ± 0.67 | 0.2329 | 0.8178 |
| Burning sensation | 1.21 ± 0.43 | 1.25 ± 0.45 | -0.2072 | 0.8376 |
| Skin reddening | 1 ± 0 | 1 ± 0 | 0 | 1 |
| Drowsiness | 2.07 ± 0.83 | 2.25 ± 0.75 | -0.5708 | 0.5735 |
| Troubled focusing | 1 ± 0 | 1 ± 0 | 0 | 1 |
| Mood swings | 1 ± 0 | 1 ± 0 | 0 | 1 |

**Table S2. Complete statistical report for the FC comparisons at the PCU.** FDR correction (q = 0.1, $n_{comparisons}=18$) yielded a critical p-value of 0.0425. Significant results are bolded. PCU: precuneus; DMN: default mode network.

|  |  | 2-sample ratio t-test | | | Repeated-measures t-test | | | | | |
| --- | --- | --- | --- | --- | --- | --- | --- | --- | --- | --- |
|  |  | *Verum* vs *Sham* | | | *Verum* | | | *Sham* | | |
|  | Areas | Cohen's d | t-stat | p-value | Cohen's d | t-stat | p-value | Cohen's d | t-stat | p-value |
| AEC-c | PCU-PCU | -1.3268 | -3.3726 | **0.0025** | -0.353 | -2.3152 | **0.0376** | 0.2575 | 2.5845 | **0.0254** |
|  | PCU-DMN | -1.4322 | -3.6406 | **0.0013** | -0.3374 | -2.7491 | **0.0166** | 0.3231 | 2.4821 | **0.0305** |
|  | PCU-AAL | -1.4227 | -3.6165 | **0.0014** | -0.3117 | -2.8251 | **0.0143** | 0.2266 | 2.2942 | **0.0425** |
| ci-PLV | PCU-PCU | 0.4566 | 1.1606 | 0.2572 | 0.0702 | 0.3223 | 0.7523 | -0.2416 | 1.2744 | 0.2288 |
|  | PCU-DMN | -0.4622 | -1.1749 | 0.2516 | -0.2939 | -1.4933 | 0.1592 | 0.1595 | -0.6441 | 0.5327 |
|  | PCU-AAL | -0.1403 | -0.3567 | 0.7244 | -0.0332 | -0.1832 | 0.8574 | 0.0604 | 0.355 | 0.7293 |

**Table S3. Complete statistical report for the links between the areas of the DMN for AEC-c.** FDR correction (q = 0.1, $n_{comparisons}=28$) yielded a critical p-value of 0.0244. Significant results are bolded. SFG: superior frontal gyrus; ACC: anterior cingulate cortex; PCC: posterior cingulate cortex; Hip: hippocampus; IPG: inferior parietal gyrus; PCU: precuneus.

|  | 2-sample ratio t-test | | |
| --- | --- | --- | --- |
|  | *Stim* vs *Sham* | | |
|  | AEC-c | | |
| Areas | Cohen's d | t-stat | p-value |
| SFG-SFG | -0.3482 | -0.885 | 0.3849 |
| SFG-Rectus | -0.4279 | -1.0876 | 0.2876 |
| SFG-ACC | -0.2699 | -0.6862 | 0.4992 |
| SFG-PCC | -0.0237 | -0.0602 | 0.9525 |
| SFG-Hip | -0.5766 | -1.4657 | 0.1557 |
| SFG-IPG | -0.2536 | -0.6446 | 0.5253 |
| SFG-PCU | -0.5732 | -1.457 | 0.1581 |
| Rectus-Rectus | -0.0517 | -0.1315 | 0.8965 |
| Rectus-ACC | -0.3582 | -0.9104 | 0.3717 |
| Rectus-PCC | -0.1161 | -0.2951 | 0.7705 |
| Rectus-Hip | -0.8434 | -2.1439 | 0.0424 |
| Rectus-IPG | -0.5172 | -1.3148 | 0.201 |
| Rectus-PCU | -1.0924 | -2.7768 | **0.0105** |
| ACC-ACC | -0.4529 | -1.1513 | 0.2609 |
| ACC-PCC | -0.3933 | -0.9998 | 0.3274 |
| ACC-Hip | -0.9447 | -2.4013 | **0.0244** |
| ACC-IPG | -0.0915 | -0.2326 | 0.8181 |
| ACC-PCU | -1.1466 | -2.9146 | **0.0076** |
| PCC-PCC | -1.2553 | -3.191 | **0.0039** |
| PCC-Hip | -0.4275 | -1.0868 | 0.2879 |
| PCC-IPG | -0.5013 | -1.2743 | 0.2148 |
| PCC-PCU | -1.1999 | -3.0501 | **0.0055** |
| Hip-Hip | -0.6826 | -1.7352 | 0.0955 |
| Hip-IPG | -0.6421 | -1.6322 | 0.1157 |
| Hip-PCU | -0.4328 | -1.1003 | 0.2821 |
| IPG-IPG | -1.0074 | -2.5607 | **0.0172** |
| IPG-PCU | -0.2576 | -0.6547 | 0.5189 |
| PCU-PCU | -1.3268 | -3.3726 | **0.0025** |

**Table S4. Complete statistical report for the links between the areas of the DMN for ci-PLV.** No comparisons survived FDR correction (q = 0.1, $n_{comparisons}=28$). SFG: superior frontal gyrus; ACC: anterior cingulate cortex; PCC: posterior cingulate cortex; Hip: hippocampus; IPG: inferior parietal gyrus; PCU: precuneus.

|  | 2-sample ratio t-test | | |
| --- | --- | --- | --- |
|  | *Stim* vs *Sham* | | |
|  | ci-PLV | | |
| Areas | Cohen's d | t-stat | p-value |
| SFG-SFG | -0.5841 | -1.4847 | 0.1506 |
| SFG-Rectus | -0.6596 | -1.6766 | 0.1066 |
| SFG-ACC | -0.7898 | -2.0075 | 0.0561 |
| SFG-PCC | 0.0056 | 0.0142 | 0.9888 |
| SFG-Hip | -0.3283 | -0.8345 | 0.4122 |
| SFG-IPG | -0.7473 | -1.8995 | 0.0696 |
| SFG-PCU | -0.5195 | -1.3207 | 0.1991 |
| Rectus-Rectus | -0.5375 | -1.3662 | 0.1845 |
| Rectus-ACC | -0.6513 | -1.6555 | 0.1108 |
| Rectus-PCC | -0.1275 | -0.324 | 0.7487 |
| Rectus-Hip | -0.7375 | -1.8746 | 0.0731 |
| Rectus-IPG | -0.9007 | -2.2894 | 0.0311 |
| Rectus-PCU | -0.5727 | -1.4558 | 0.1584 |
| ACC-ACC | -0.784 | -1.9929 | 0.0578 |
| ACC-PCC | -0.2482 | -0.6309 | 0.5341 |
| ACC-Hip | -0.8251 | -2.0973 | 0.0467 |
| ACC-IPG | -0.992 | -2.5215 | 0.0187 |
| ACC-PCU | -0.6407 | -1.6287 | 0.1164 |
| PCC-PCC | 0.375 | 0.9533 | 0.35 |
| PCC-Hip | -0.2255 | -0.5731 | 0.5719 |
| PCC-IPG | -0.1894 | -0.4816 | 0.6345 |
| PCC-PCU | 0.1362 | 0.3463 | 0.7321 |
| Hip-Hip | -0.6111 | -1.5533 | 0.1334 |
| Hip-IPG | -0.6471 | -1.6449 | 0.113 |
| Hip-PCU | -0.1577 | -0.4009 | 0.692 |
| IPG-IPG | -0.5876 | -1.4936 | 0.1483 |
| IPG-PCU | -0.2485 | -0.6316 | 0.5336 |
| PCU-PCU | 0.4566 | 1.1606 | 0.2572 |
